# ProtFinder: finding subcellular locations of proteins using protein interaction networks

**DOI:** 10.1101/2022.01.11.475836

**Authors:** Aayush Grover, Laurent Gatto

**Affiliations:** International Institute of Information Technology Bangalore, Bengaluru, India; Computational Biology and Bioinformatics Unit, de Duve Institute, Université catholique de Louvain, Brussels, Belgium

**Keywords:** Subcellular Location, Protein Interaction Networks, Deep Learning, Multi-label Classifier

## Abstract

Protein subcellular localization prediction plays a crucial role in improving our understanding of different diseases and consequently assists in building drug targeting and drug development pipelines. Proteins are known to co-exist at multiple subcellular locations which make the task of prediction extremely challenging. A protein interaction network is a graph that captures interactions between different proteins. It is safe to assume that if two proteins are interacting, they must share some subcellular locations. With this regard, we propose ProtFinder – the first deep learning-based model that exclusively relies on protein interaction networks to predict the multiple subcellular locations of proteins. We also integrate biological priors like the cellular component of Gene Ontology to make ProtFinder a more biology-aware intelligent system. ProtFinder is trained and tested using the STRING and BioPlex databases whereas the annotations of proteins are obtained from the Human Protein Atlas. Our model obtained an AUC-ROC score of 90.00% and an MCC score of 83.42% on a held-out set of proteins. We also apply ProtFinder to annotate proteins that currently do not have confident location annotations. We observe that ProtFinder is able to confirm some of these unreliable location annotations, while in some cases complementing the existing databases with novel location annotations. The source code for ProtFinder is available at https://github.com/UCLouvain-CBIO/ProtFinder.

## 1 Introduction

Protein participates in every process of the cell, from metabolic reactions to immune response. The functional properties of a protein are known to strongly correlate with the spatial information of the protein within the cell. Moreover, studying subcellular localisation of proteins can help in drug discovery, and in gene annotation [1, 2]. For instance, the proteins that tend to exist near the membrane of the cell could act as target sites for the drug being trialed.

Traditional methods in biology to learn the localisation of proteins are time and resource consuming processes. Hence, recent works have focused on computational methods to accurately predict the protein subcellular localisation. Particularly, the deep learning pipelines are now widely used to complement the experimental approaches. While these pipelines rely on experimental validation, they can be used to add scalability to the robustness of experimentation. Some of the earlier computational methods treated this task as a multi-class single-label classification [3, 4, 5]. However, with recent advancements it was learnt that a protein can either co-exist in multiple locations within a cell or move around different subcellular components [6]. This multi-label assumption makes the task of prediction significantly more arduous. A few recent works [2, 7] build multi-label classifiers for this task using the amino-acid sequence of proteins. Wan et. al [2] builds an SVM-based classifier to predict across 10 subcellular locations whereas Cheng et. al [7] uses Gene Ontology (GO) information and the Gaussian kernel regression model to identify the localisation across 14 locations. More recently, the focus has shifted to predict the subcellular locations of protein through image-based feature extraction [8, 9]. Liu et. al [8] uses multi-view image features from microscopy images. These features are used to predict protein localization across 7 subcellular location. A multi-label classifier using stacked Autoencoders [10] and Random Forest [11] is proposed. Along the similar lines, Su et. al [9] uses deep Convolutional Neural Networks (CNNs) [12] to extract features from microscopy images of proteins and uses these features to predict across 7 subcellular locations through an SVM [13] classifier.

Another important source of information for this task is the protein interaction networks. A protein interaction network is essentially a graph with proteins as nodes and and interactions as edges. The extent of interaction is treated as the weight of the edge in the network. If two proteins interact with each other, then they are likely to share some subcellular locations. Integrating such biological priors will assist in building more explainable and biologically relevant classifiers. Moreover, these networks can also be used to annotate unlabelled proteins within the network. In 2019, Pan et. al [5] developed a deep learning (RNN) single-label classifier called node2loc and used the protein interaction networks to classify across 16 subcellular locations.

Despite the progress so far, there is still a scope of improvement. There is a need to use more biological priors to build task-relevant estimators. It is also important to build multi-label classifiers because single-label classifiers have very limited usability. The rapid improvement in the availability of databases like STRING [14], BioPlex [15], and Human Protein Atlas (HPA) [16] allow for more deep learning approaches to be built for this task.

In this work, we treat this task of protein subcellular localisation prediction as a multi-class multi-label classification task while using biological priors like the protein interaction networks and the cellular component (CC) of Gene Ontology (GO). We propose ProtFinder, a multi-label deep neural model that predicts the localisation of the protein across 28 subcellular locations. Motivated by node2loc, we use a graph learning algorithm to first obtain the latent representation of the network. This representation is then passed through a deep multi-label classifier that outputs the log likelihood of the given protein to reside in each of the 28 different locations.

The structure of rest of the article is as follows: The datasets used and their processing is explained in Section 2. Section 3 discusses the ProtFinder model and its implementation details. The metrics used, model performance, and some inferences are drawn in Section 4. Finally, the concluding remarks and future directions of this work are discussed in Section 5.

## 2 Materials

In this section, we will first discuss the datasets used in Section 2.1. This will be followed by understanding their processing steps in Section 2.2.

### 2.1 Data Sources

We combine information from various sources of datasets – STRING [14] and BioPlex [15] in order to build a more rich and comprehensive protein interaction network. The proteins in the network were annotated using the Human Protein Atlas (HPA) [16]. In the protein interaction network datasets, each row represents an interaction which consists of a pair of protein ID and a numerical measure of interaction whereas, in the HPA, each row consists of a protein ID, it’s possible locations, and the reliability of this localisation information. The reliability is classified into “Enhanced”, “Supported”, “Approved”, and “Unreliable” with “Enhanced” being the most reliable and ‘Unreliable” being the least reliable.

The HPA consisted of location annotations of 12,390 proteins across 32 subcellular locations. The STRING database holds information of 5,879,727 interactions across 19,354 unique proteins. On the other hand, the BioPlex database has information regarding the 118,162 interactions between 13,689 unique proteins.

### 2.2 Preprocessing

The proteins obtained from the STRING and BioPlex databases were first mapped with the dataset from the HPA. The protein IDs are mapped across different databases using the online tool of Ensembl BioMart [17]. While combining the HPA data with the protein interaction data, we do not consider “Unreliable”, “Supported” and “Approved” annotations as valid annotations as we want to train our model on the most reliable location annotations. Therefore, we only map the locations with reliability class “Enhanced”.

After annotating the STRING and BioPlex databases with the HPA database, we obtain interactions where (1) both proteins have location annotations, (2) exactly one protein has location annotation, and (3) neither proteins have location annotations. The interactions belonging to the last category were removed from the combined database. This was done to ensure that we do not end up with a graph with sparse annotations. Moreover, the BioPlex database also has a measure of unreliability of an interaction. The measure of interaction in the BioPlex database was a value in the range [0, 1] whereas, it was in the range [0, 999] in the STRING database. Therefore, the value was normalized to [0, 1] in the STRING database to maintain uniformity. In the BioPlex dataset, those interactions that have the probability of being wrong greater than 5% were dropped. This would warrant only those interactions that are sure to exist in reality. After removing these interactions, we were left with 3,126,817 interactions from the STRING database and 73,664 interactions from the BioPlex database which when combined forms a network consisting of 3,186,865 weighted edges (interactions) and 23,165 nodes (proteins).

We then observe the location distribution across the STRING and BioPlex datasets as shown in Figure 1. This figure depicts the count of proteins belonging to each subcellular location (Table 1) in the HPA. The 4 subcellular locations – “Rods & Rings”, “Aggresome”, “Microtubule ends”, and “Cleavage furrow” were not considered in our task as these locations did not have sufficient samples in STRING as well as BioPlex. Therefore, our analysis is restricted to 28 subcellular locations.

**Figure 1:**
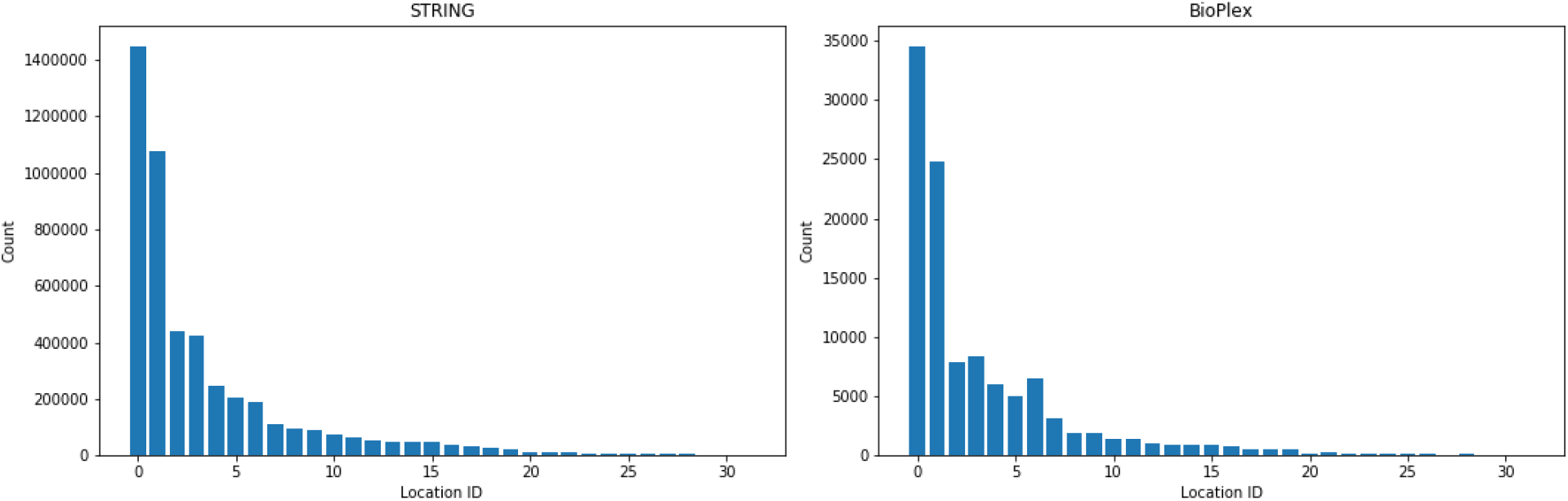
Frequency of proteins with a particular location label.

**Table 1:** Subcellular locations considered in our analysis and their respective identifiers. Location ID corresponds to the identifiers used in preprocessing whereas Class ID corresponds to each of the binary classifier of ProtFinder.

| Subcellular Location | Location ID | Class ID |
| --- | --- | --- |
| Nucleoplasm | 0 | 25 |
| Cytosol | 1 | 7 |
| Plasma membrane | 2 | 27 |
| Vesicles | 3 | 28 |
| Mitochondria | 4 | 18 |
| Golgi apparatus | 5 | 11 |
| Nucleoli | 6 | 23 |
| Endoplasmic reticulum | 7 | 8 |
| Nuclear bodies | 8 | 20 |
| Nuclear speckles | 9 | 22 |
| Cell Junctions | 10 | 2 |
| Centrosome | 11 | 4 |
| Nuclear membrane | 12 | 21 |
| Microtubules | 13 | 15 |
| Nucleoli fibrillar center | 14 | 24 |
| Actin filaments | 15 | 1 |
| Intermediate filaments | 16 | 12 |
| Centriolar satellite | 17 | 3 |
| Focal adhesion sites | 18 | 10 |
| Cytokinetic bridge | 19 | 5 |
| Cytoplasmic bodies | 20 | 6 |
| Mitotic spindle | 21 | 19 |
| Midbody | 22 | 16 |
| Lipid droplets | 23 | 13 |
| Lysosomes | 24 | 14 |
| Rods & Rings | 25 | - |
| Endosomes | 26 | 9 |
| Midbody ring | 27 | 17 |
| Peroxisomes | 28 | 26 |
| Aggresome | 29 | - |
| Microtubule ends | 30 | - |
| Cleavage furrow | 31 | - |

After combining the HPA data with the protein interaction datasets, we obtain a dataset with location information (along with reliability of the location annotations) of both the proteins interacting as well as the measure of interaction. Due to the assumption that proteins that are interacting must be interacting at a common location, we redefine the location annotations of each protein. This is done as shown in Algorithm 1.

This algorithm allows us to remove those locations that might be rarely occurring in reality or a mis-annotation in the HPA. We call such annotations “noisy”. The hyperparameter *α* is defined as the de-noising threshold which helps in filtering and eventually, removing the noisy location annotations. The range of *α* is [0,1] where *α* = 0 implies that all annotations are noisy whereas *α* =1 implies that no annotation is noisy. The optimal value of *α* is learnt by studying Figures 2 and 3. These figures help us understand the average number of locations that any random protein can co-exist in. It can be observed that for *α* = 0.6, we obtain the location count distribution across the network that matches with the HPA database (Figure 3). We can now be sure that the number of locations occupied by any given protein in our processed combined dataset is in agreement with the experimental dataset from HPA database. These new annotations will be used as the ground truth values to train ProtFinder.

**Figure 2:**
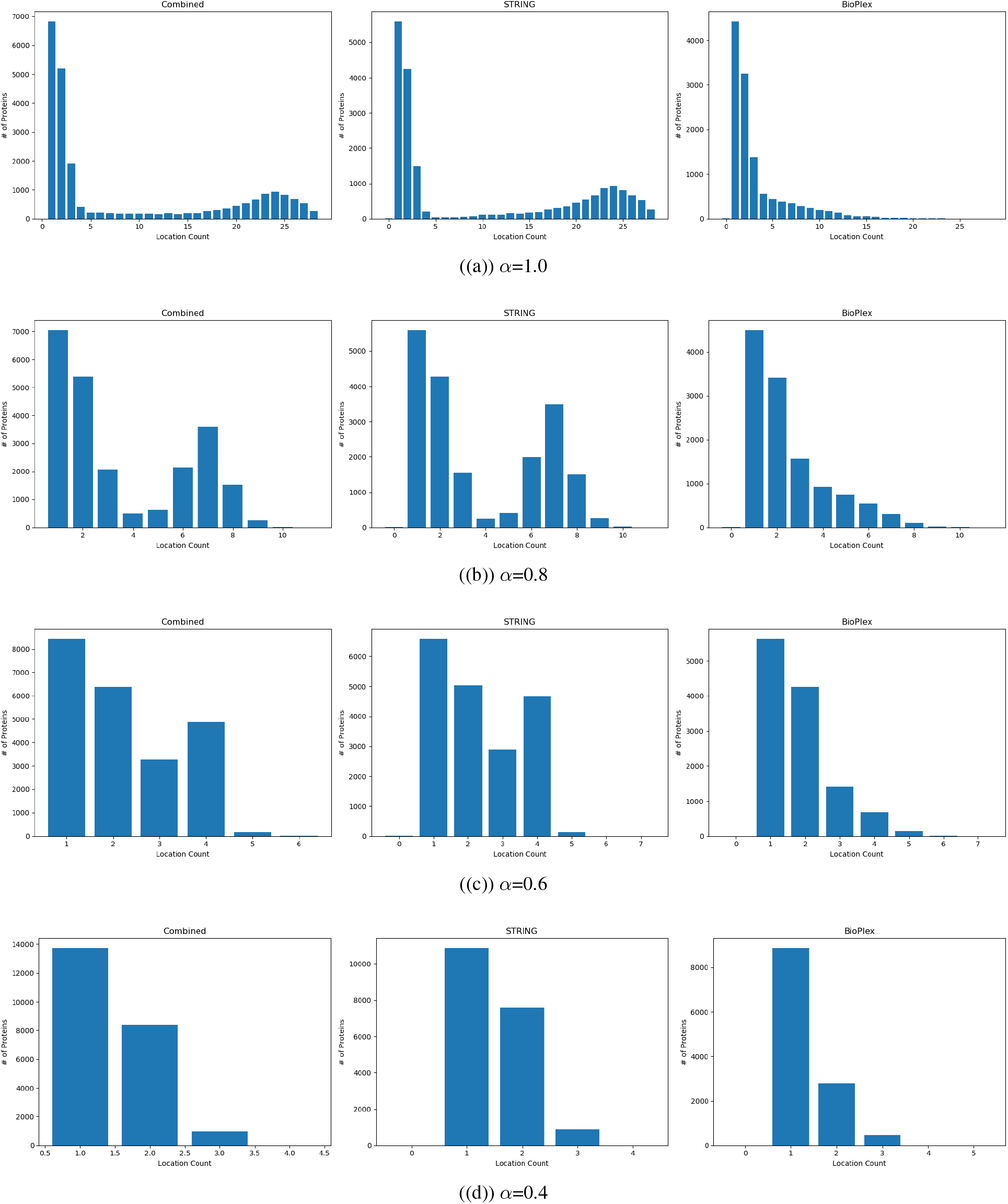
Distribution of number of locations a protein is localised to for various de-noise thresholds.

**Figure 3:**
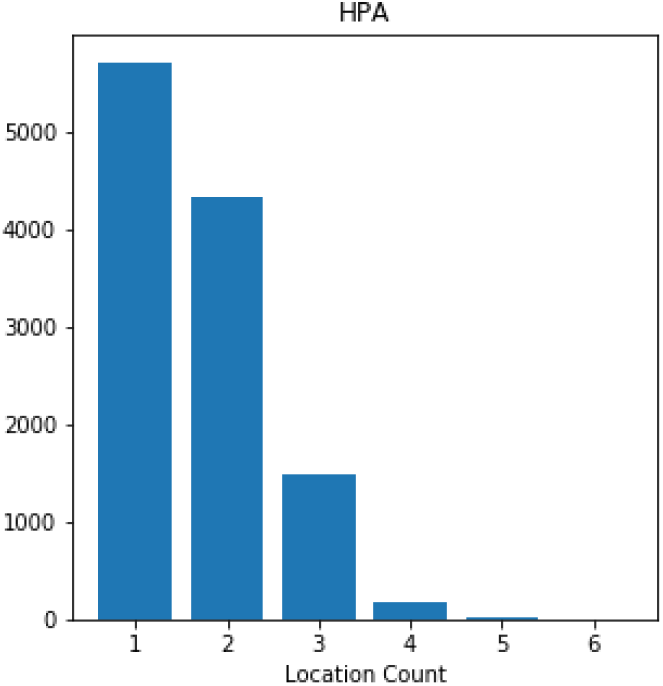
Distribution of number of locations a protein is localised to in the HPA database.

#### Algorithm 1 Re-annotations of locations of proteins

~~~
**Require:** Interaction network (*G*) = List(*interaction*) where a given *interaction* = (Protein1, Location1, Reliability1 Protein2, Location2, Reliability2, Weight).
 Locations and reliabilities are obtained from HPA whereas Weight is the measure of interaction.
**Require:** *α* – De-noising threshold
 **for all** *interaction* **in** *G* **do**
  *location*1 = interaction[Location1]
  *location*2 = interaction[Location2]
  **if** *location*1 = NULL and *location*2 = NULL **then**
    Remove *interaction*
  **else if** *location*1 = NULL **then**
    *interaction*[Locations] = *location*2
  **else if** *location*2 = NULL **then**
    *interaction*[Locations] = *location*1
  **else**
    *interaction*[Locations] = *location*1 ⋂ *location*2
  **end if
 end for
 for all** *protein* **in** *G* **do**
    *count* = dict()
    *count* [all locations] = 0
    **for all** *interaction* **in** *G* **where** *interaction*[Protein1] = *protein* **or** *interaction*[Protein2] = *protein* **do
      for all** *location* in interaction[Locations] **do**
        *count*[*location*] = *count*[*location*] + 1
     **end for
  end for**
  Sort *count* by value (computed above)
  Normalize the values in *count* to range [0, 1]
  Retain maximum locations with cumulative sum of *count* values ≤ *α*
  new_location for *protein* = retained locations (above)
**end for**
~~~

Upon following the above steps, we acquire a protein interaction network where some proteins have location annotations whereas others do not. Precisely, the protein interaction network has 9,639 unannotated proteins which are kept aside for drawing inferences. The remaining 13,523 annotated proteins are used for training and validating the ProtFinder model (80% for training and 20% for validation). We ensure that the train and validation datasets follow a similar distribution across different subcellular locations. We observe a spearman’s rho of 97.21% with a p-value of 6.86 × 10^−18^ between the train and validation distributions. This validation data will be called the test data until Section 4.3.

## 3 Methods

After combining and processing datasets from various sources, in this section, we define our ProtFinder model and discuss its implementation details. The overview of the ProtFinder model can be found in Figure 4. The ProtFinder model can be divided into two modules -

- **Representation Learning** - An unsupervised representation learning module that learns that latent representation of each protein (node embeddings) through the protein interaction network.
- **Multi-label Classifier** - A supervised deep neural multi-label classifier that predicts the likelihood of a protein to exist in each of the 28 subcellular locations.

**Figure 4:**
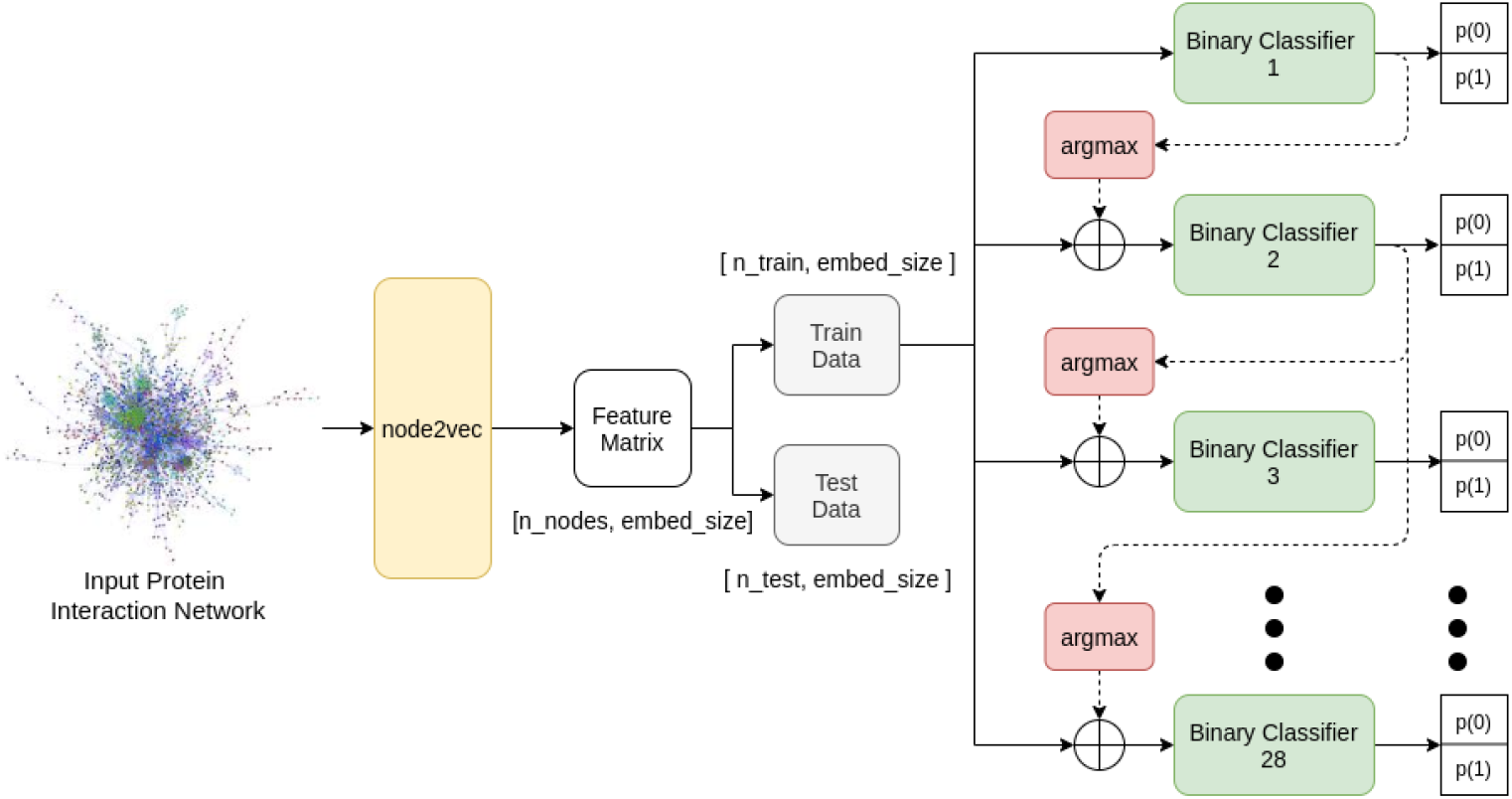
Overview of ProtFinder

Section 3.1 explains the Representation Learning module in detail and explains the algorithms used. The Multi-label Classifier is defined in Section 3.2. Section 3.2 also describes how various biological information can be integrated into the Multi-label Classifier. Finally, the implementation details are explained in Section 3.3.

### 3.1 Representation Learning

Representation Learning is a paradigm of machine learning and deep learning that aims to represent information in a concise and effective manner using supervised or unsupervised learning algorithms. A better representation method can make learning of the downstream tasks easier. Representation learning methods have been widely used across various fields – Natural Language [18], Image Analysis [19], and Speech [20]. Network Representation Learning [21] is a sub-field of Representation Learning which focuses on learning representations of a network. A representation of a network is considered better than the others if it can preserve the graph topology while also clustering similar nodes together in the embedding space.

In ProtFinder, the Representation Learning module is of the type Network Representation Learning as it aims to generate node embeddings for the input protein interaction network. Node embeddings are a continuous-valued vector representation of a given node in a network. We use a graph learning algorithm called node2vec [22] to generate the embeddings of size 128. Each protein (annotated and unannotated) in the network is now represented by a continuous-valued 128-dimensional vector. Node2vec uses a two step process to learn the node embeddings – (1) uses random walk sampling strategy to first obtain a chain of nodes (or proteins) (2) applies the word2vec [23] algorithm to learn the embeddings of each node in the chain. This process is repeated multiple times. We also try some of the more recent methods to obtain node embeddings – AttentionWalk [24] and GraphSAGE [25]. However, the performance of ProtFinder was best with embeddings generated using node2vec. The comparison between the high-dimensional embeddings generated by Node2vec and AttentionWalk is depicted in Figure 5. This is done by first mapping the 128 dimensional space to a 2 dimensional space using first the PCA [26] and then the t-SNE [27] algorithm. For node2vec, the configuration was – *n_components:* 0.9, *perplexity:* 75, *early_exaggeration:* 12, *random_state:* 10, and *n_iter:* 700 whereas for AttentionWalk, the configuration was – *n_components:* 0.99, *perplexity:* 62, *early_exaggeration:* 12, *random_state:* 10, and *n_iter:* 1000. These configurations were chosen as these gave the most separable results for each of these embedding generation algorithms.

**Figure 5:**
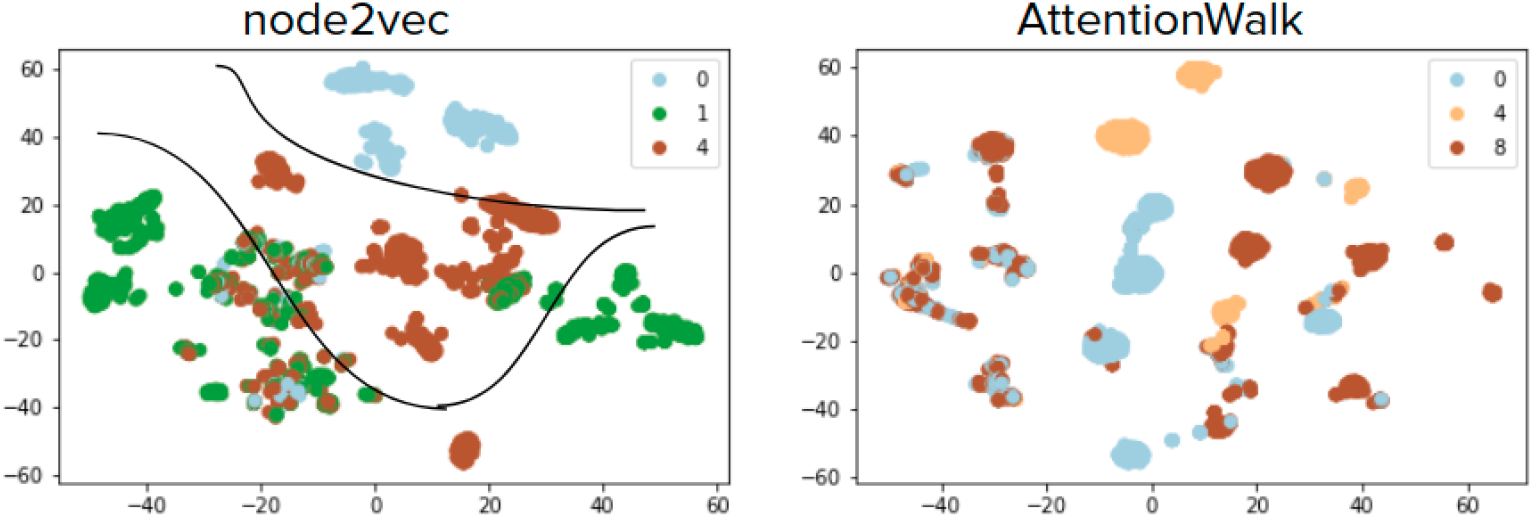
Comparison between node embeddings generated by node2vec (left) and AttentionWalk (right) using t-SNE plots. The embeddings are generated for the proteins that belong to the 3 most commonly occurring locations – Plasma membrane, Cytosol, Nucleoplasm. The tickers in node2vec are 0:Plasma membrane, 1:Nucleoplasm, 4:Cytosol and in AttentionWalk are 0:Cytosol, 4:Plasma membrane, 8:Nucleoplasm

Clearly, the embeddings generated by node2vec are more separable than those generated by AttentionWalk. Hence, we use node2vec to generate the embeddings of each protein. The generated embeddings are stored in a matrix where each row represents a protein. Therefore, we obtain a matrix of size = number of proteins × embedding size (i.e. 23162 × 128). This embedding matrix is used to train and evaluate our deep learning-based multi-label classifier.

### 3.2 Multi-label Classifier

The task of protein subcellular localisation prediction is that of a multi-label classification because any given protein can co-exist at multiple locations within the cell. Therefore, we need a solution that, when given a protein embedding, predicts all the possible locations of the protein at once. In this regard, we propose a multi-label classifier as described in Figure 4.

The proposed multi-label classifier consists of 28 binary classifiers (*BC*_1_, *BC*_2_,…, *BC*_28_) – each binary classifier *BC_i_* corresponding one location (Class ID *i* from Table 1). Motivated by Pan et. al [5], binary classifier *BC_i_* consists of an LSTM cell [28] which learns the hidden patterns within the generated protein embeddings. This is followed by a fully-connected layer which creates non-linear combinations of the learnt hidden patterns. The output of the fully connected layer is represented by a vector *z* of length *K* where *K* is the number of classes for each classifier. Since, each classifier is a binary classifier, *K* = 2. The fully-connected layer is followed by the softmax activation function (Equation 1) which is used to output the probability of the given protein to exist at subcellular location *i*. The overview of each binary classifier is given in Figure 6.

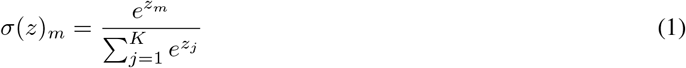

**Figure 6:**
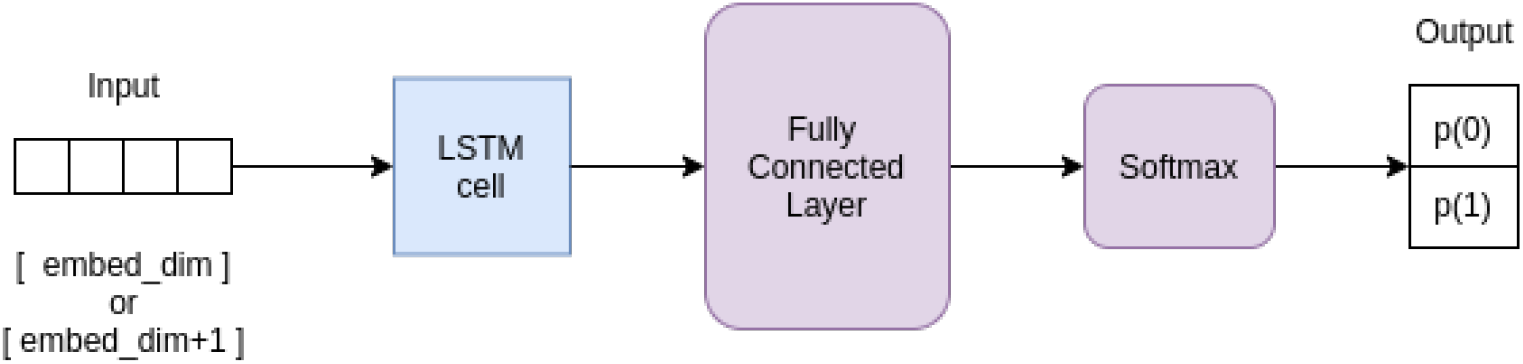
Architecture of each binary classifier in ProtFinder

Using the obtained probability, we then use a threshold value *t_i_* to binarize the output i.e. the output of binary classifier *BC_i_* is 1 if the output probability is greater than *t_i_* and the output is 0 otherwise (Equation 2). Through this, the model predicts the multiple possible subcellular locations of a give protein (all locations i such that output of *BC_i_* is 1).

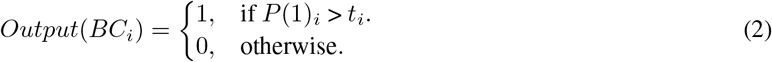

The embedding matrix obtained through the Representation Learning module (Section 3.1) is split into training and test proteins. Training proteins are the 13,523 annotated proteins whereas the remaining 9,639 unannotated proteins are used to test the ProtFinder model. Each binary classifier is trained using the training proteins. Binary Cross-Entropy (BCE) loss is used to train each binary classifier and is given by Equation 3. Here, *y_i_* is 1 if output of *BC_i_* is 1 and *ŷ_i_* is 1 if location *i* is in the annotation of the given protein.

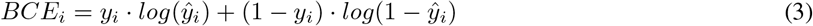

#### 3.2.1 Relation Between Subcellular Locations

It is important to note that all locations mentioned in Table 1 are not independent of each other. They indeed have relations between them which are described by the Cellular Components of Gene Ontology (GO-CC) [29]. The GO-CC describes locations at the subcellular and macro-molecular levels. A relation is said to exist from subcellular location *C_i_* to subcellular location *C_j_* if there is a path from *C_i_* to *C_j_* such that level of *C_i_* in GO-CC is greater than level of *Cj* in GO-CC and none of the remaining 26 subcellular locations lie in this path. A schematic of this relation construction is given in Figure 7. In Figure 7(a), the nodes that are of interest (locations mentioned in Table 1) are labelled in purple. The relations can be stored in the form of a disconnected directed acyclic graph (DAG) where each node is a subcellular location and each directed edge depicts the relation between two subcellular locations (Figure 7(b)). The edge-list of this DAG is extracted from the GOView tool of WebGestalt [30] can be found in Table 2. The subcellular locations that are not mentioned in Table 2 are independent subgraphs in the DAG.

**Figure 7:**
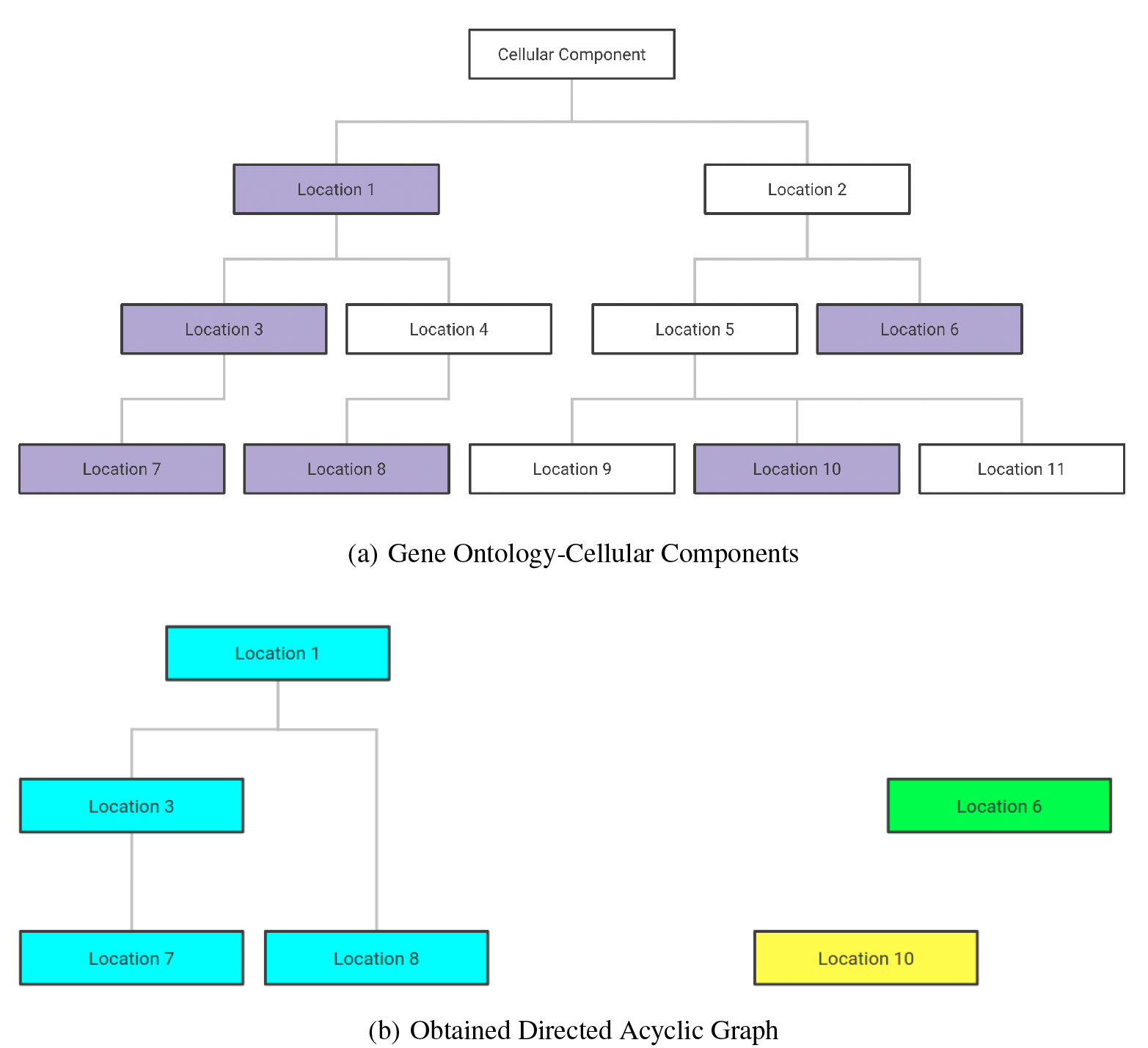
(a) Complete graph of relations given by GO-CC. The locations marked in purple are locations of interest. (b) The obtained DAG graph from the locations of interest. Each disconnected subgraph is denoted in a different colour. **NOTE**: The data for this image is hypothetical. This is a toy example.

**Table 2:**
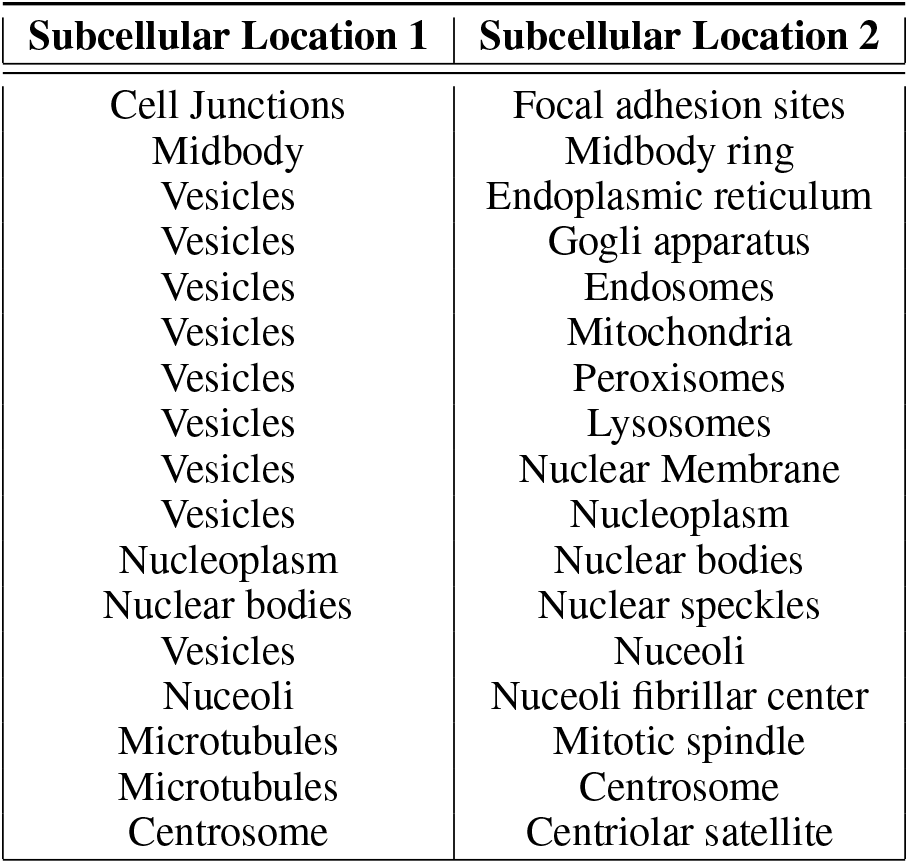
Edge-list of the DAG created from the GOView tool. Each relation is from Subcellular Location 1 to Subcellular Location 2.

| Subcellular Location 1 | Subcellular Location 2 |
| --- | --- |
| Cell Junctions | Focal adhesion sites |
| Midbody | Midbody ring |
| Vesicles | Endoplasmic reticulum |
| Vesicles | Gogli apparatus |
| Vesicles | Endosomes |
| Vesicles | Mitochondria |
| Vesicles | Peroxisomes |
| Vesicles | Lysosomes |
| Vesicles | Nuclear Membrane |
| Vesicles | Nucleoplasm |
| Nucleoplasm | Nuclear bodies |
| Nuclear bodies | Nuclear speckles |
| Vesicles | Nuceoli |
| Nuceoli | Nuceoli fibrillar center |
| Microtubules | Mitotic spindle |
| Microtubules | Centrosome |
| Centrosome | Centriolar satellite |

It is important to integrate this information about relation between different subcellular locations within the model. Without loss of generality, let us assume that a directed relation exists between Class ID 1 (*C*_1_) and Class ID 8 (*C*_8_) such that *C*_1_ → *C*_8_ (as shown in Figure 7). Therefore, output of *BC*_1_ can influence the result of *BC*_8_. To capture this influence, the output of *BC*_1_ is binarized using threshold *t*_1_ and this is then added to the node embeddings used to classify for subcellular location *C*_8_. This gives us a new binary feature for prediction of subcellular location *C*_8_. We now use (embedding size +1) input features for *BC*_8_ to predict the probability of the given protein to exist at subcellular location 8. These relations are denoted by dotted lines in Figure 4.

#### 3.2.2 Defining Thresholds

We initially run the experiments using each threshold *t_i_* = 0.5 ∀*i* ∈ [1, 28]. However, each class ID can have different threshold values and we can learn the class thresholds to improve the performance of ProtFinder. To further examine this, we observed the distance between predictions and target of each class in the training dataset. The predictions range from 0 to 1 whereas targets are either 0 or 1. Therefore, all the values of (target-predictions) between −0.5 and 0.5 are correctly classified whereas values in range [0.5, 1] represent False Negatives and values in the range [−1, −0.5] represent False Positives. As we can see in Figure 8, there are several cases of misclassifications for each class ID that can be avoided. For example, in class ID 1, we can increase the threshold value so that the misclassifications around −0.6 can now be correctly classified. Similarly, we can improve the model performance by decreasing the threshold *t*_12_. Therefore, decide the thresholds for each class ID independently can help build a more accurate multi-label classifier.

**Figure 8:**
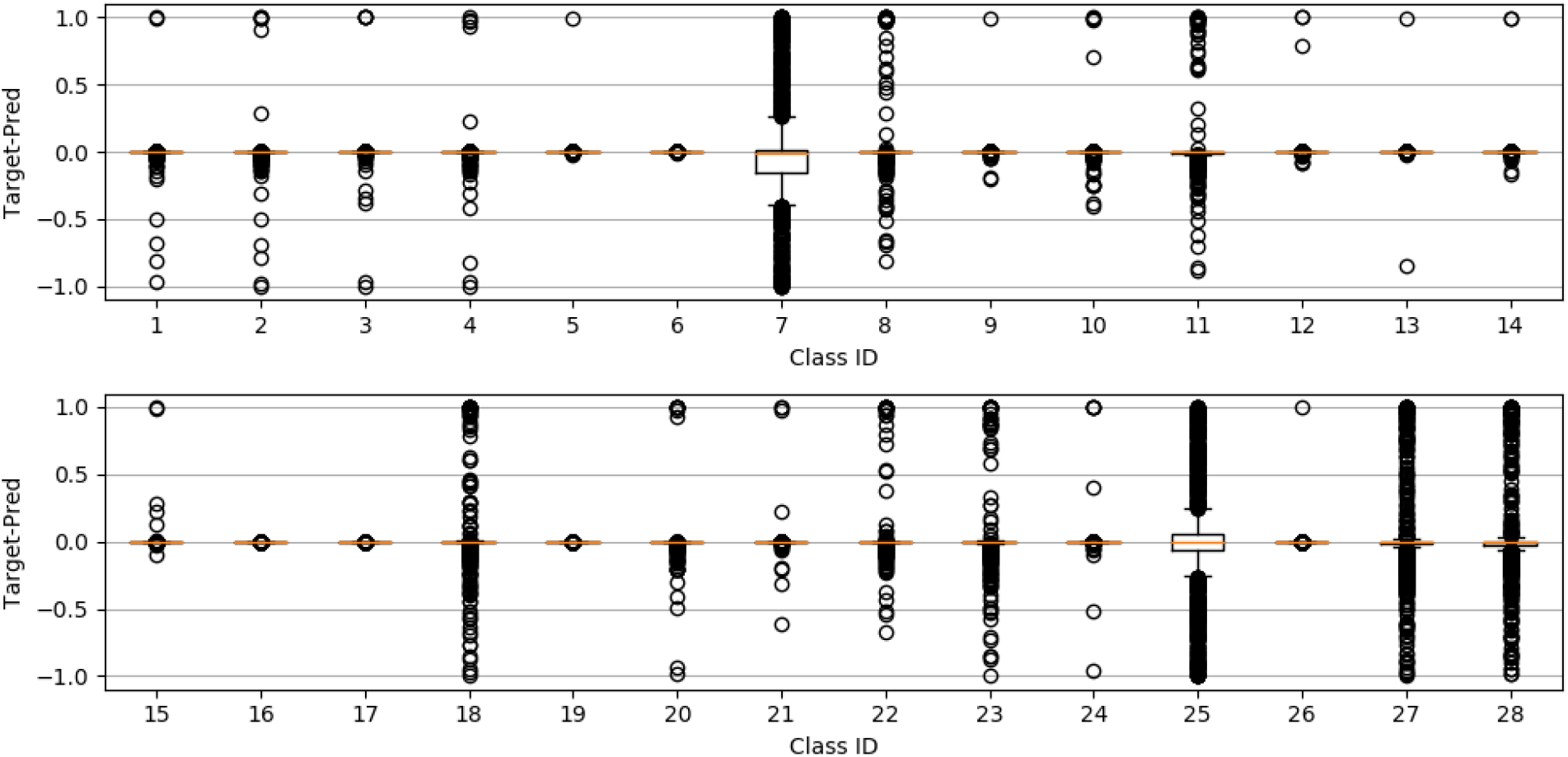
Box Plot showing (Target-Prediction) values for each class ID

### 3.3 Implementation Details

Manual computation of each of the 28 threshold values can be time-consuming and might result in inaccurate thresholds due to human errors. Therefore, we treat each of the 28 thresholds as hyperparameters. Along with these, we also keep training epoch as a hyperparameter. The optimal values of these 29 hyperparameters are computed using the Ax^1^ hyperparameter optimizer which uses the bayesian optimization algorithm. The Ax algorithm was trained for 5 trials as each trial is computationally expensive. The 5 trials were run on a 64-core machine with 256GB DDR3 ECC RAM for approximately 24 days. The final set of hyperparameters can be found in Table 3.

**Table 3:** Final set of hyperparameters as obtained by Ax

| Hyperparameter | Value | Hyperparameter | Value |
| --- | --- | --- | --- |
| epochs | 8 | $t_{15}$ | 0.4693 |
| $t_1$ | 0.5156 | $t_{16}$ | 0.5088 |
| $t_2$ | 0.5840 | $t_{17}$ | 0.5048 |
| $t_3$ | 0.5057 | $t_{18}$ | 0.5986 |
| $t_4$ | 0.5743 | $t_{19}$ | 0.5002 |
| $t_5$ | 0.5021 | $t_{20}$ | 0.5795 |
| $t_6$ | 0.5054 | $t_{21}$ | 0.5241 |
| $t_7$ | 0.5318 | $t_{22}$ | 0.4758 |
| $t_8$ | 0.4802 | $t_{23}$ | 0.5410 |
| $t_9$ | 0.4809 | $t_{24}$ | 0.5396 |
| $t_{10}$ | 0.5200 | $t_{25}$ | 0.5683 |
| $t_{11}$ | 0.4852 | $t_{26}$ | 0.5075 |
| $t_{12}$ | 0.3569 | $t_{27}$ | 0.5524 |
| $t_{13}$ | 0.5033 | $t_{28}$ | 0.4645 |
| $t_{14}$ | 0.5090 | | |

The Representation Learning module was used as is from node2vec [22]. The Multi-label Classifier was implemented using PyTorch [31] library implemented for Python. All source codes can be found on https://github.com/UCLouvain-CBIO/ProtFinder.

## 4 Experiments & Results

In this section, we first define the metrics used to test the performance of ProtFinder in Section 4.1. Section 4.2 compares the performance of different versions of ProtFinder. Finally, we draw some inferences from the trained ProtFinder model in Section 4.3.

### 4.1 Metrics

To test the performance of ProtFinder, we look at the following metrics –

- Accuracy [32] is the percentage of correctly classified instances (Equation 4).
- Recall or sensitivity [33], which denotes the fraction of positive instances that the model was able to retrieve (Equation 5).
- Specificity [33], which denotes the fraction of negative instances that the model was able to correctly classify (Equation 6).
- The area under receiver operating characteristics (AUC-ROC) [34], which is the area under the curve plotted when the true positive rate is plotted against the false positive rate.
- Matthews correlation coefficient (MCC) [35], which is a metric that takes into consideration all the cells of the confusion matrix (Equation 7).

The above-mentioned metrics are averaged across all binary classifiers. This will help in understanding the overall performance of the multi-label classifier. Due to the significant class imbalance within each binary classifier, MCC is the best metric to test the model performance as this metric is agnostic of the label statistics.

In the following expressions, TP, TN, FP and FN denote true positive, true negative, false positive and false negative, respectively.

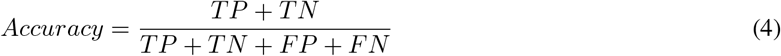

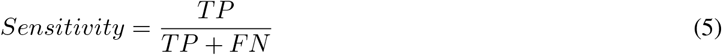

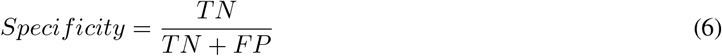

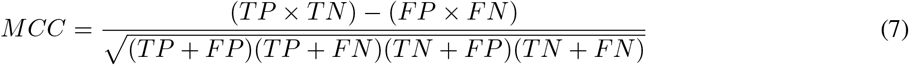

### 4.2 Model Performance

The choice of LSTM cell in each binary classifier is motivated by Pan et. al [5]. However, it is not justified. It is also necessary to observe the effect of DAG (Section 3.2.1) on the overall model performance. We empirically compare the performance of ProtFinder where each binary classifier contains the LSTM cell (ProtFinder) against ProtFinder where each binary classifier only consists of fully connected layers (ProtFinder-FCC) and ProtFinder where the binary classifiers are treated independent of each other (ProtFinder-noDAG). The metrics defined in Section 4.1 are used to test the performance of the model on the validation set of proteins. The comparison between ProtFinder, ProtFinder-noDAG and ProtFinder-FCC, and the performance of each of these methods can be found in Table 4. It is important to note that the performances of ProtFinder, ProtFinder-noDAG and ProtFinder-FCC are computed on their respective set of best hyperparameters obtained after 5 trials of Ax.

**Table 4:** Performance of ProtFinder-FCC vs ProtFinder-noDAG vs ProtFinder. The best performances are marked in bold.

| Model | Accuracy | Sensitivity | Specificity | AUC-ROC | MCC |
| --- | --- | --- | --- | --- | --- |
| ProtFinder-FCC | 0.9825 | 0.9927 | 0.7492 | 0.8765 | 0.7745 |
| ProtFinder-noDAG | 0.9870 | 0.9952 | **0.7975** | **0.9013** | 0.8318 |
| ProtFinder | **0.9872** | **0.9957** | 0.7943 | 0.9000 | **0.8342** |

As observed in Table 4, the ProtFinder outperforms ProtFinder-FCC in all the metrics (only marginally on some metrics but significantly on the others) whereas giving a comparable performance to ProtFinder-noDAG. Crucially, ProtFinder shows an improvement of approximately 5% over ProtFinder-FCC on the MCC metric and 0.25% over ProtFinder-noDAG. ProtFinder-noDAG outperforms ProtFinder in specificity and AUR-ROC by approximately 0.3% and 0.15% respectively. Therefore, the LSTM-based ProtFinder models can be considered to be the best-performing classifiers. However, we believe that ProtFinder (with biological prior as DAG) is a better model as it integrates biological information into the deep learning model while giving a slightly better performance on the MCC metric. We also observe class-wise scores – average accuracy score for each binary classifier of the ProtFinder model in Table 5. The accuracies are computed over proteins in the validation dataset.

**Table 5:** Average performance of each subcellular location on the test data along with the distribution of proteins in the training and test data. Class ID corresponds to each of the binary classifier of ProtFinder. Table is sorted in the ascending order of MCC. The table is divided between classifiers that make random predictions and the classifiers that make non-random predictions.

| Subcellular Location | Class ID | Accuracy (test) | MCC (test) | Number of Proteins (train) | Number of Proteins (test) |
| --- | --- | --- | --- | --- | --- |
| Midbody ring | 17 | 0.9996 | nan | 1 | 0 |
| Mitotic spindle | 19 | 0.9996 | nan | 3 | 0 |
| Nucleoli fibrillar center | 24 | 0.9959 | nan | 38 | 10 |
| Midbody | 16 | 0.9993 | -0.0074 | 2 | 2 |
| Cytoplasmic bodies | 6 | 0.9985 | -0.0074 | 11 | 2 |
| Lipid droplets | 13 | 0.9985 | -0.0068 | 8 | 3 |
| Lysosomes | 14 | 0.9981 | -0.0068 | 9 | 3 |
| Endosomes | 9 | 0.9981 | -0.0063 | 8 | 4 |
| Peroxisomes | 26 | 0.9985 | -0.0063 | 14 | 1 |
| Cytokinetic bridge | 5 | 0.9985 | -0.0063 | 21 | 4 |
| Actin filaments | 1 | 0.9959 | -0.0060 | 42 | 7 |
| Centriolar satellite | 3 | 0.9970 | -0.0059 | 12 | 5 |
| Centrosome | 4 | 0.9926 | 0.1704 | 85 | 21 |
| Microtubules | 15 | 0.9918 | 0.2299 | 69 | 18 |
| Nuclear bodies | 20 | 0.9885 | 0.2750 | 86 | 19 |
| Cell Junctions | 2 | 0.9907 | 0.3444 | 81 | 14 |
| Intermediate filaments | 12 | 0.9937 | 0.3537 | 67 | 20 |
| Focal adhesion sites | 10 | 0.9937 | 0.3627 | 47 | 12 |
| Golgi apparatus | 11 | 0.9777 | 0.4477 | 329 | 79 |
| Nuclear membrane | 21 | 0.9955 | 0.4601 | 50 | 14 |
| Endoplasmic reticulum | 8 | 0.9836 | 0.5910 | 245 | 61 |
| Nucleoli | 23 | 0.9810 | 0.6260 | 302 | 78 |
| Nuclear speckles | 22 | 0.9855 | 0.6361 | 190 | 48 |
| Cytosol | 7 | 0.9174 | 0.7821 | 2937 | 723 |
| Plasma membrane | 27 | 0.9632 | 0.8018 | 1117 | 272 |
| Vesicles | 28 | 0.9602 | 0.8146 | 1071 | 274 |
| Mitochondria | 18 | 0.9773 | 0.8288 | 706 | 180 |
| Nucleoplasm | 25 | 0.9472 | 0.8905 | 5217 | 1300 |

Through Table 5, we can observe that most binary classifiers predict the outcomes with high accuracy (greater than or equal to 95%). The only outliers being binary classifiers for subcellular locations Cytosol and Nucleoplasm. Nonetheless, even for these classes, the classifier accuracies are greater than 90% (91.74% and 94.72% respectively). We also observe that MCC score for many subcellular locations is close to 0 (upper half of Table 5) indicating that the model is using random guess to predict these locations. To understand why this would be happening, we compare the distribution of MCC scores with the distribution of proteins used to train and test the binary classifiers (Table 5). We obtain a spearman’s rho of 92.41% with a p-value of 4.34 × 10^−11^ (between MCC and train data) and spearman’s rho of 93.18% with a p-value of 1.32 × 10^−11^ (between MCC and test data). This shows that the binary classifiers of those subcellular locations that do not hold many proteins, do not perform well on MCC. This observation is in agreement with the deep learning theory which states that it is difficult to train a deep neural model with very few samples. Upon further analysis, we observe that most of the subcellular locations in the upper half of Table 5 are locations that do not have many proteins in the protein interaction networks (STRING and BioPlex). Some of these subcellular locations do not have reliable annotations in the HPA. However, this is also true for other subcellular locations for which the binary classifiers perform well. Therefore, we can conclude that adding more protein interaction data would help improve the classification performance of proteins into these subcellular locations.

Multi-class multi-label learning models is often susceptible to predicting outputs that have a different distribution as compared to the target. Figure 9 depicts the distribution of number of subcellular locations predicted by the ProtFinder as compared to the distribution of number of subcellular locations of proteins in the annotations of the test data. We can observe that the output predictions of the ProtFinder model tends to follow a similar distribution to the target distribution, exceptions being ProtFinder predicting no subcellular locations for 349 proteins. Apart from just observing the distributions, we also compute the performance of ProtFinder for each number of subcellular locations predicted. Table 6 shows the performance of ProtFinder based on number of subcellular locations predicted for proteins in the test data. We observe no MCC value when the model predicts no location because in this case true positive (TP) and false positive (FP) are both 0 thereby making the MCC value undefined.

**Figure 9:**
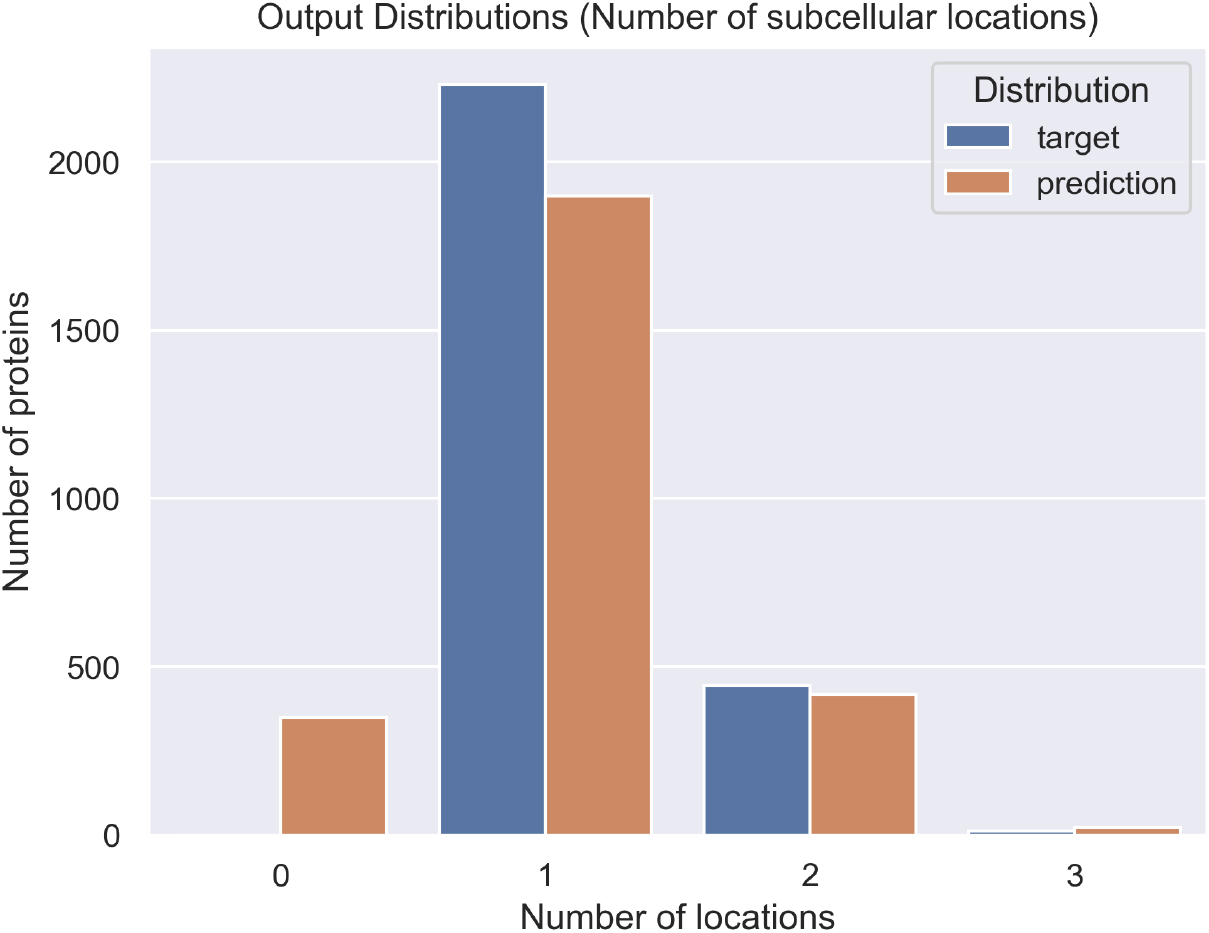
Comparison of the distribution of number of locations in the target and predicted subcellular locations. The comparison is based on the predictions on the test data.

**Table 6:** MCC scores of ProtFinder based on different number of subcellular locations predicted in the test data.

| Number of Locations | MCC |
| --- | --- |
| 0 | - |
| 1 | 0.9121 |
| 2 | 0.8620 |
| 3 | 0.7702 |

Being the first method to use protein interaction networks for multi-class multi-label classification of protein subcellular locations, ProtFinder cannot be compared with existing multi-label classifiers that use amino-acid sequence as input. However, we compare the performance of ProtFinder with node2loc [5]. Node2loc aims to solve a simpler task of predicting a single subcellular location out of 16 available locations with an MCC score of 0.812. ProtFinder, on the other hand, is able to solve a more challenging task of multi-class prediction of localisation of proteins across 28 locations with an average MCC score of 0.8342 (Table 4). This improvement in performance for a significantly challenging task makes ProtFinder the new state-of-the-art for this task.

### 4.3 Inferences

The main objective of ProtFinder model was to predict the subcellular locations of unannotated proteins (proteins that do not have reliable annotations in the Human Protein Atlas). In this section, we show the predictive performance on ProtFinder on these proteins. It is difficult to predict the performance of ProtFinder on those proteins that do not have any concrete information regarding their subcellular locations. Therefore, we cherry-pick some proteins that have unreliable location information. Through this, we can confirm the unreliable locations. It will, however, be difficult to draw inferences from instances where there is disagreement between the predicted locations and unreliably known locations. We try to inspect the overlap in these cases with the UniProt database [36]. Nonetheless, in order to draw any final conclusions, we would require experimental validation.

#### 4.3.1 Case Study 1: Proteins with Agreement

Here, we would discuss case study of some proteins with similar annotations by ProtFinder and by the “Unreliable” class of locations in the Human Protein Atlas (HPA).

According to the Human Protein Atlas (HPA), the Dynein light chain LC8-type 2 protein (ENSP00000477310) belongs to subcellular locations Nucleoplasm, Nuclear Bodies, and Cytosol whereas, according to the ProtFinder predictions, it belongs to subcellular locations – Nucleoplasm (probability=99.63%), Nuclear Bodies (probability=90.12%), Cytosol (probability=89.2%), Midbody (probability=81.95%), and Cell Junctions (probability=79.36%). However, according to UniProt, this protein exists in Cytosol, Nucleus and Plasma Membrane.

Similarly, for the Mitochondrial ribosomal protein L15 (ENSP00000260102) is majorly known to be present in the Mitochondria (UniProt and HPA) and this is in agreement with the prediction of ProtFinder – Mitochondria (probability=99.97%), and Mitotic Spindle (probability=70.58%).

Finally, another positive case of predictions by ProtFinder is for the EEF1A lysine methyltransferase 1 protein (ENSP00000372202). The HPA labels Cytosol and Plasma Membrane as the subcellular locations of this protein. The UniProt labels this protein to be in Cytosol. The ProtFinder model predicts Cytosol (probability=92.22%). The second highest probability of a subcellular location is that of Plasma Membrane with 29.20%.

The above three examples show how ProtFinder can be used to confirm the uncertain annotations of some proteins while also trying to identify other possible subcellular locations.

### 4.4 Case Study 2: Proteins with Disagreement

We were also able to identify a few cases where the prediction from ProtFinder was in disagreement with the not-so-reliable classes of HPA. The conflicting locations could be a misclassification by ProtFinder or could be a highly unreliable annotation in HPA.

For instance, according to the HPA, ADP ribosylation factor 6 protein (ENSP00000298316) majorly exists in Cytosol whereas ProtFinder predicts this particular protein to exist in Plasma Membrane (probability=94.82%), Vesicles (probability=84.41%), Peroxisomes (probability=81.62%), Lipid Droplets (probability=76.12%), Midbody Ring (71.86%), Cytoplasmic Bodies (probability=71.56%), and finally Cytosol (69.03%). However, UniProt assigns this protein to Cytosol, Plasma Membrane, Golgi Apparatus, Endosomes, and Focal Adhesion Cites. Therefore, it is not clear if this particular protein exists only in Cytosol, Plasma Membrane or a subset of the subcellular locations predicted by ProtFinder.

Similarly, Proteasome subunit beta 4 (ENSP00000290541) exists in Nucleoplasm and Mitochondria according to the HPA. However, the ProtFinder model predicts Focal Adhesion Sites, Cytosol, Lysosomes, Lipid Droplets, Endosomes, Centriolar Satellite, and Mitotic Spindle (all > 90%) with a higher probability than Nucleoplasm (89.50%) and Mitochondria (5.19%) are not in the top predictions. However, this protein exists in Nucleoplasm, Cytosol, and Mitochondria according to UniProt. Therefore, there is no common consensus about the location of this protein.

The above examples show how in some predictions the unreliable locations from the HPA don’t completely match with the predictions of ProtFinder. This might diminish the practical usage of ProtFinder. Some of the disagreements between HPA and ProtFinder are confirmed by the UniProt. However, we might require clinical experimentation to confirm if ProtFinder is indeed able to give better predictions than the unreliable annotations of the HPA.

## 5 Conclusion

ProtFinder is a multi-class multi-label classifier that accurately annotates subcellular locations of a given protein. Unlike most of the existing approaches that use the amino-acid sequence or microscopy imaging to predict subcellular locations, ProtFinder relies solely on the protein interaction networks. This is based on the assumption that if two proteins are interacting, they are likely to share some subcellular locations. However, with the increasing research in multi-modal learning techniques, integrating the three different modes of data – amino-acid sequence, microscopy imaging, and protein interaction networks, would be a promising direction in building comprehensive systems to predict protein subcellular locations. Node2loc [5] also uses protein interaction networks to predict subcellular locations of the proteins. However, it predicts a single location for a given protein which is not the case in reality [6]. Proteins can co-exist in multiple locations or move from location to another, thereby making it necessary to predict its localisation. To this extent, ProtFinder predict the protein localisation across multiple subcellular locations which is more complex and challenging.

The ProtFinder model can classify a protein accurately across 28 subcellular location making ProtFinder the most widely usable method (to the best of our knowledge) for the protein subcellular localization prediction task. The task of multi-label prediction is a challenging task with significant class imbalance. Despite this, ProtFinder predicted the subcellular locations of unseen proteins with an average MCC score of 83.42% making it a robust and reliable model. ProtFinder also uses the Cellular Components of Gene Ontology to incorporate the relation between different subcellular locations. This integrates the biological prior that is used by ProtFinder to improve its predictions. The model is also able to capture the distribution of number of locations to good extent making it a practically usable approach. We also observe that ProtFinder performs better for some subcellular locations as compared to others. This can be mitigated by enriching the protein interaction network databases like STRING and BioPlex which will consequently reduce the class imbalance.

The ProtFinder model firstly uses a graph learning algorithm called node2vec to generate protein embeddings. It would be interesting to see the impact of graphical neural networks like Message Passing Neural Networks (MPNNs) [37] to generate these embeddings. The generated embeddings are then used by the LSTM-based multi-class multi-label classifier to accurately predict the subcellular locations of the proteins. Such deep neural embedding generation methods are better feature extractors and can be used to make the ProtFinder model end-to-end trainable. This means that the embeddings will incorporate the information about the localisation of proteins making the embeddings rich of information for this task.

We also show that the ProtFinder model can be used to predict subcellular localisation of proteins that are not annotated with a high level of confidence in the Human Protein Atlas. The model predictions can be used to strengthen the confidence of existing annotations or to conflict the unreliable predictions by providing better annotations in the existing databases. This is also shown with the help of UniProt. However, the conflicts need to be resolved by wet-lab experimentation. Nonetheless, based on the performance of ProtFinder on the validation set and some of the inferences drawn in Section 4, we can confidently utilize this method to predict subcellular locations of proteins. Such a deep learning-based method with the assistance of biological priors can help provide better relevant solutions for many medical tasks. Specifically, ProtFinder can play a major role in annotating proteins which can further be used to aid the drug targeting and drug discovery processes.

## Footnotes

1 https://ax.dev/

## Notes

### Competing Interest Statement

The authors have declared no competing interest.

### Summary of Updates

Added URL to software GitHub repo in abstract and section 3.3.

